# Intraflagellar Transport Selectivity Occurs with the Proximal Portion of the Trypanosome Flagellum

**DOI:** 10.1101/2024.12.13.628094

**Authors:** Aline Araujo Alves, Jamin Jung, Gaël Moneron, Humbeline Vaucelle, Cécile Fort, Johanna Buisson, Cataldo Schietroma, Philippe Bastin

**Affiliations:** Trypanosome Cell Biology Unit, Institut Pasteur, Université de Paris Cité, INSERM U1201, Paris, France; Institut Pasteur, Paris, France; Abbelight, www.abbelight.com, Cachan, France

## Abstract

Intraflagellar transport (IFT) trains move bidirectionally along the doublet microtubules (DMTs) of the axoneme within the flagellum. In *Trypanosoma brucei*, IFT trains predominantly associate with four of the nine DMTs. Using high-resolution microscopy, we reveal how this selective association is put in place. IFT proteins form a ring surrounding the 9 DMTs on top of the transition fibres. Volume electron microscopy revealed densities along all DMTs in the proximal portion of the flagellum, exhibiting thinner, shorter profiles with branches absent in mature IFT trains. As the axoneme extends within the flagellar pocket, IFT trains are detected but are often positioned outside DMTs 3-4/7-8. After the axoneme exits the flagellar pocket, IFT trains localise exclusively to DMTs 3-4 and 7-8. Super-resolution and expansion microscopy demonstrated that IFT proteins follow the same distribution as the IFT-like densities. This suggests they represent IFT trains undergoing assembly and/or disassembly and reveals their unexpected ability to shift from one DMT to another.

**Summary:** In *Trypanosoma brucei*, intraflagellar transport (IFT) trains selectively associate with specific axonemal microtubules. Using advanced microscopy, this study reveals how this restriction occurs at the proximal portion of the flagellum during the assembly and/or disassembly of IFT trains.

## Introduction

Sensory and motile cilia are microtubule-based organelles constructed by a specialised transport mechanism called intraflagellar transport (IFT). IFT consists of multiprotein complexes assembled into polymers, or IFT trains, that move bidirectionally along the axonemal microtubules (Kozminski et al., 1993). Specific molecular motors drive the IFT trains: kinesin-2 powers the anterograde trains, while cytoplasmic dynein 2 carries the retrograde trains (Kozminski et al., 1995; Pazour et al., 1998; Porter et al., 1999). IFT proteins and molecular motors are distributed along the length of the cilia but are highly concentrated around the ciliary base, where the assembly and disassembly of IFT trains take place (Buisson et al., 2013; Hibbard et al., 2021; Mijalkovic et al., 2017; Prevo et al., 2015; Wingfield et al., 2017).

IFT is essential for the building of the core of the cilium, the axoneme, which is composed of nine outer doublet microtubules (DMTs) arranged in a cylinder with or without a central pair of singlet microtubules. The cilium is constructed from the basal body and is separated from the cytoplasm by a ciliary gate, which functions as a barrier between these two environments. This gate begins with the distal appendages, known as transition fibres, which connect the distal portion of the basal body to the ciliary membrane (Anderson, 1972). Distal to the transition fibres, the transition zone appears, also composed of nine outer DMTs but lacking the central pair. The DMTs of the transition zone are connected to the ciliary membrane by Y-links (Gilula and Satir, 1972), which are present along its entire length, creating a size-exclusion filter for protein diffusion (Breslow et al., 2013; Kee et al., 2012; Lin et al., 2013). Due to this barrier, IFT is essential for the entry of ciliary proteins that cannot diffuse through the ciliary gate, making it crucial for cilia assembly (Craft et al., 2015).

IFT trains are long polymers composed of periodic repeats of two multimeric complexes, IFT-A and IFT-B, together with the IFT motors and the cargoes (Jordan et al., 2018; Pigino et al., 2009). These components are sequentially assembled at the ciliary base to form the anterograde trains. This process starts with a core of IFT-B, to which IFT-A is added, followed by the association of the cargoes, and lastly, the IFT kinesin (Wingfield et al., 2017). In the green alga *Chlamydomonas reinhardtii*, IFT trains assemble into linear arrays hanging from the cytosol, passing through the transition fibres, and tethering at the proximal portion of the transition zone (van den Hoek et al., 2022). IFT trains moving along the transition zone are also observed in the cilia of chemosensory neurons of *C. elegans* (Mitra et al., 2024). By contrast, retrograde trains are less electron-dense and harder to detect by structural biology (Stepanek and Pigino, 2016; van den Hoek et al., 2022).

*Trypanosoma brucei*, the causative agent of African trypanosomiasis, possesses a motile cilium, or flagellum, that is essential for parasite survival (Kohl et al., 2003). The trypanosome flagellum contains a canonical 9+2 axoneme and features an additional extra-axonemal structure known as the paraflagellar rod (PFR), which extends alongside the axoneme (Vickerman, 1962). The PFR is a lattice-like filamentous structure attached to the axoneme, forming connections with the flagellar membrane and the plasma membrane through the flagellum attachment zone (FAZ). This connection maintains the DMT 7 consistently facing the cell body (Farina et al., 1986; Hughes et al., 2017; Lacomble et al., 2009). Trypanosomes possess a flagellar pocket, an invagination of the plasma membrane from where the flagellum emerges from the cell body (Halliday et al., 2021). At the proximal portion of the flagellum within the flagellar pocket, trypanosomes exhibit a transition zone of approximately 300 nm in length (Trépout et al., 2018), followed by a segment of the axoneme that is free of the PFR. The PFR only attaches to the DMTs 4-7 of the axoneme after the flagellum exits the flagellar pocket.

Trypanosomes possess all 22 *IFT* genes (Morga and Bastin, 2013). These proteins have been shown to traffic along the flagellum and are essential for its construction (Absalon et al., 2008). Electron microscopy observations revealed that IFT trains are exclusively associated with DMTs 3-4 and 7-8 in procyclic-form trypanosomes (Bertiaux et al., 2018). Since these two sets of DMTs are on opposite sides of the axoneme, high-resolution live microscopy could reveal both anterograde and retrograde transport on each track. However, the mechanisms involved in the targeting of specific microtubules for IFT in trypanosomes remain unknown.

How are only four out of nine DMTs selected by IFT trains? This selectivity may originate at the level of the flagellum base, where the IFT basal pool is localised. One hypothesis proposes that IFT proteins concentrate exclusively at the base of the two sets of microtubules where IFT trains are observed. Alternatively, regardless of the basal pool surrounding all microtubules, the assembly of IFT trains could occur specifically on the microtubules where IFT trains are present (Bertiaux et al., 2018; Mallet and Bastin, 2019).

To understand how IFT selectively targets four DMTs in trypanosomes, we used multiple high-resolution techniques to map the distribution of IFT at the proximal end of the flagellum. We found that the IFT basal pool exhibits a ring-like aspect encircling all DMTs at the level of the transition fibres and the proximal transition zone. Within the transition zone and the axoneme inside the flagellar pocket, densities distribute along multiple DMTs. These densities resemble IFT trains but are thinner, shorter, and display branching not seen in mature trains. At the region of the axoneme where the flagellum exits the flagellar pocket, both these densities and typical IFT trains – thicker, longer, and denser – are present and often located on most DMTs, including outside the two tracks. As the flagellum extends beyond the pocket and the PFR associates with DMTs 4-7, IFT trains become restricted to two distinct tracks. The IFT-B protein IFT172 shares the same distribution as the IFT-like densities, suggesting they represent IFT trains undergoing assembly and/or disassembly. Our findings highlight the proximal flagellum as critical for IFT selectivity in trypanosomes and reveal the unexpected ability of trains undergoing assembly (or disassembly) to shift from one DMT to another.

## Results

### IFT basal pool surrounds all doublet microtubules of the transition zone

To investigate the hypothesis that IFT selectivity for a set of DMTs starts at the base of the flagellum, we characterised the IFT basal pool. We used *T. brucei* procyclic-form cells, the life cycle stage present in the midgut of the insect vector, expressing mCh::RP2, a protein localised to the transition fibres (Stephan et al., 2007), stained with an antibody against mCherry and a monoclonal antibody against IFT172 (Absalon et al., 2008). Observations were made using stochastic optical reconstruction microscopy (STORM) and direct optical nanoscopy with axial localised detection (DONALD) (**Fig. 1**). In the cell shown in **Fig. 1 A**, which has two flagella, two parallel lines are observed for IFT172 in the proximal region of the growing flagellum (**Fig. 1 A**, arrowheads). These lines likely correspond to the two sets of DMTs of the axoneme where IFT trains are known to be present in trypanosomes (Bertiaux et al., 2018). The concentration of IFT172 at the flagellum base is evident, and its signal partially overlaps with RP2 but is positioned more apically (**Fig. 1 A**). A side view of the transition zone region (**Fig. 1 B**) shows the IFT pool (green) distal to the transition fibres (magenta), with a region of overlap. Therefore, the IFT pool is situated partially above the transition fibres in the proximal portion of the transition zone. Additionally, thin lines of the IFT172 signal are detected along the distal portion of the transition zone, possibly indicating IFT trains under construction and/or disassembly (**Fig. 1 B**, arrowhead). These images are consistent with observations of IFT material progression during train construction in *C. elegans* (Prevo et al., 2015), RPE-1 cells (Yang et al., 2019) and *Chlamydomonas reinhardtii* (van den Hoek et al., 2022).

**Figure 1:**
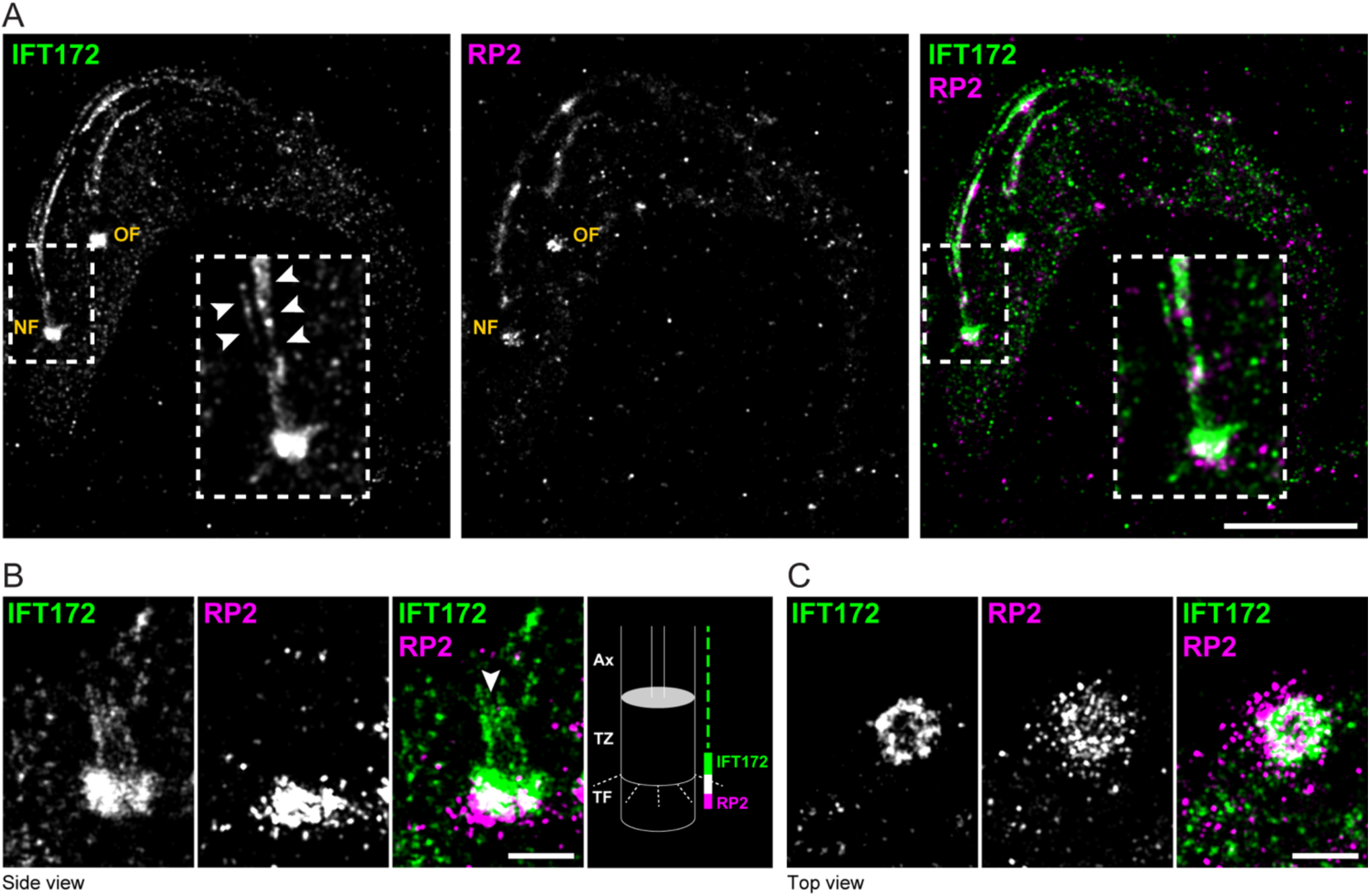
The IFT basal pool localises at the proximal transition zone in the direct vicinity of the transition fibre marker RP2. The localisation of IFT172 was examined using direct optical nanoscopy with axially localised detection on fixed procyclic-form trypanosomes expressing mCh::RP2 using antibodies against IFT172 (green) and mCherry (magenta) as indicated. **(A)** View of a whole biflagellate cell. The inset on the left panel is a magnified view of the proximal region of the new flagellum (NF), showing the distribution of IFT trains on both sides of the axoneme (arrowheads). The same inset in the right panel provides a closer look at the base of the flagellum, highlighting the proximity of IFT172 and mCh::RP2. NF = new flagellum, OF = old flagellum. **(B)** The side view image of the transition zone region indicates that the IFT basal pool is located more distally than the transition fibre protein RP2. The merged portion in white illustrates the partial overlap. The arrowhead indicates possible IFT trains under construction. The right panel provides a schematic representation of RP2 and IFT172 distribution relative to the flagellar components. TF = transition fibres, TZ = transition zone, Ax = axoneme. **(C)** Representative top view image of the transition zone region, showing IFT172 arranged in a ring-like structure. Scale bars: 500 nm.

A top view of the flagellum base (**Fig. 1 C**) reveals that IFT172 is arranged in a ring-like aspect. This pattern was confirmed in 16 analysed 3D STORM stacks, with the IFT172 ring diameter measured at 356 ± 26 nm (*n* = 16), closely matching the flagellar membrane diameter surrounding the transition zone as determined by scanning transmission electron microscopy (STEM) (352 ± 27 nm) (Trépout et al., 2018). This is consistent with the presence of the IFT pool as a ring around the 9 DMTs in the proximal portion of the transition zone, overlapping with the transition fibres. Thus, we can reject the hypothesis that the IFT basal pool is only present at the base of the DMTs 3-4 and 7-8.

### IFT trains localise to DMTs 3-4 and 7-8 in bloodstream-form trypanosomes

Another hypothesis posits that IFT polymerisation occurs exclusively at DMTs 3-4 and 7-8. To test this, we examined the base of the flagellum, where IFT assembly occurs. We used focused ion beam scanning electron microscopy (FIB-SEM) to precisely position IFT trains within flagella volume. FIB-SEM offers sufficient resolution to distinguish between the DMTs while enabling the monitoring of long flagellar segments, thereby allowing the 3D observation of IFT distribution (Bertiaux et al., 2018). Because bloodstream-form trypanosomes, the life cycle stage present in the mammalian host, are thinner than their procyclic counterparts, they were chosen as a model, thus increasing the number of flagella available within a given volume. As IFT has not been investigated in this life cycle stage, we first aimed to describe it. We used bloodstream-form cells expressing mNG::IFT81 to image IFT in living cells (**Fig. S1 A** & **Video 1**). Temporal projections of the IFT81 signal revealed a concentration at the flagellum base and its proximal portion, followed by an even distribution of IFT material along the flagellar length. Kymographs showed particles moving in both anterograde and retrograde directions (**Fig. S1 A**), exhibiting profiles similar to those observed in procyclic form trypanosomes (Buisson et al., 2013).

Subsequently, we investigated the position of IFT trains using transmission electron microscopy (TEM), where they appear as electron-dense particles located between the axonemal microtubules and the flagellar membrane (Absalon et al., 2008; Kozminski et al., 1995; Pigino et al., 2009). The DMTs were numbered clockwise relative to the orientation of the dynein arms, with DMT 1 positioned opposite to the PFR and DMT 7 identified by its longer connector to the PFR (**Fig. S1 B**)(Zhang et al., 2021). Among the 60 IFT trains observed in 41 flagella (**Fig. S1 C**), more than 90% were positioned on DMTs 3-4 and 7-8, with 31.7% located on DMTs 3-4 and 60% on DMTs 7-8 (**Fig. S1 D**). These findings confirm that IFT trains predominantly travel along two tracks in bloodstream-form trypanosomes, similar to previous observations in the procyclic stage (Bertiaux et al., 2018).

Given that IFT distribution in bloodstream-form trypanosomes is comparable to previous observations, we characterised it along the flagellum length using FIB-SEM (**Fig. 2**). For this analysis, bloodstream-form trypanosomes were fixed, embedded in epoxy resin, and imaged by FIB-SEM on a slice-and-view mode, which generated serial images of a large sample volume (Vanwalleghem et al., 2015). IFT trains were positioned as before based on the axoneme’s orientation relative to the PFR and the cell body, with DMTs 7-8 facing the side of the flagellum attached to the cell body (**Fig. 2 A**) (Hughes et al., 2017). The long DMT7 connection with the PFR could also be distinguished (**Fig. 2 A-C**, magenta asterisk) (Sherwin and Gull, 1989). Since the central pair (CP) of microtubules does not rotate (Branche et al., 2006; Gadelha et al., 2006; Ralston et al., 2006), it could also serve as a reference; DMTs 3-4 and 7-8 are always positioned perpendicularly to the CP. In the FIB-SEM volume, IFT trains were identified as densities between the axoneme and the flagellar membrane, present in three or more consecutive slices and continuous alongside the microtubules (**Fig. 2 B-D**). The generated FIB-SEM series included 29 flagella from 25 cells (19 cells with one flagellum and six cells with two flagella, from which two flagella were too short to be reconstructed). These flagella and their associated IFT trains were segmented and reconstructed in different colours (**Fig. 2 E**, left panel; **Video 2**). A single reconstructed flagellum shows the IFT trains (**Fig. 2 E**, right panel, arrowheads; **Video 3**) distributed on opposite sides of the axoneme corresponding to DMTs 3-4 and 7-8. Following the reconstruction of all IFT trains obtained within the volume using the criteria described before, 224 IFT trains were positioned into pairs of neighbouring DMTs (**Fig. 2 F**), as IFT trains are often positioned between two DMTs on electron microscopy observations. We found that more than 94% of the IFT trains were positioned on DMTs 3-4/7-8, with 54.5% on DMTs 3-4 and 39.7% on DMTs 7-8, consistent with the TEM results.

**Figure 2:**
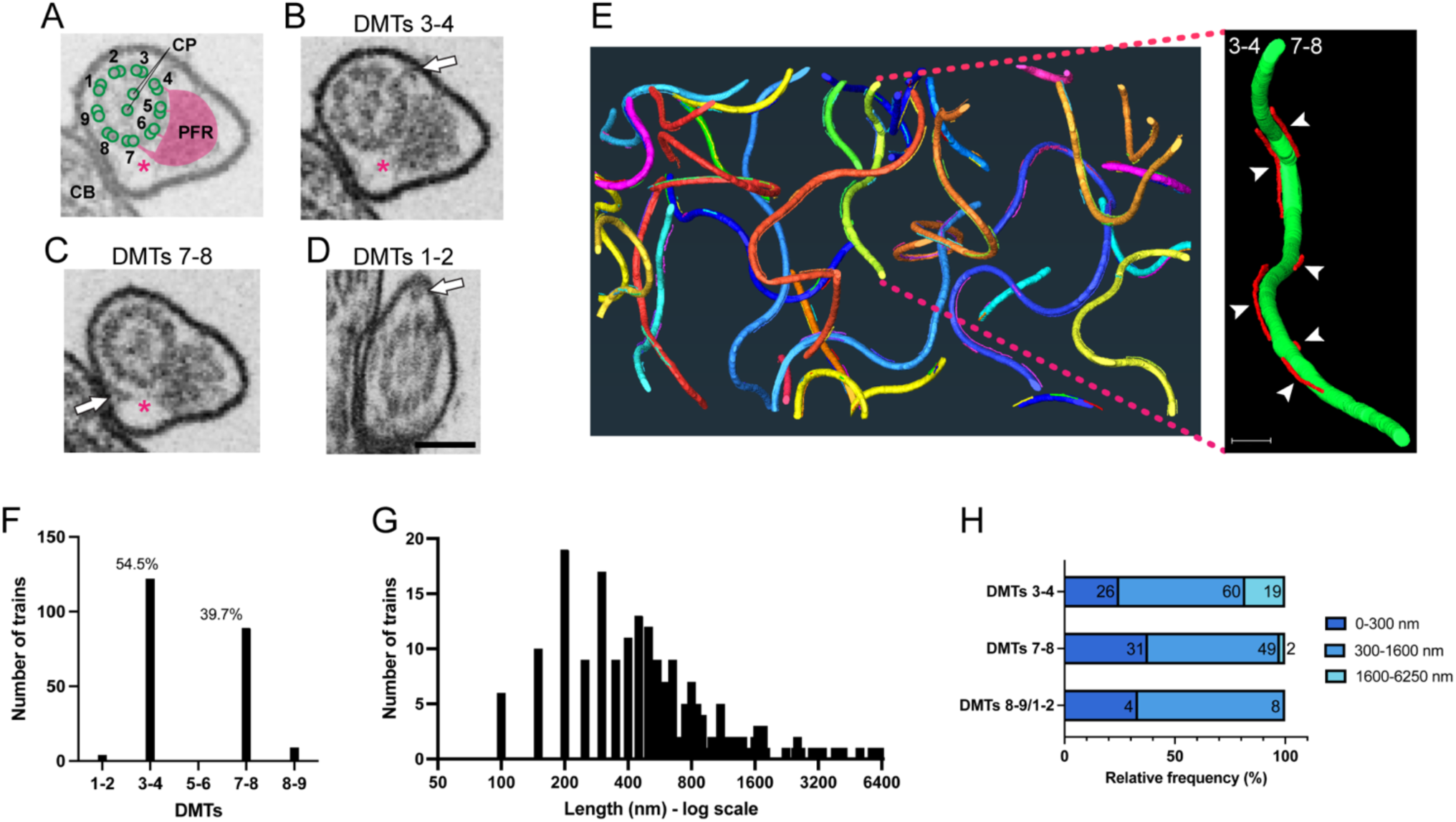
IFT trains distribute into four axoneme doublet microtubules in the bloodstream form of *Trypanosoma brucei*. Flagella and IFT trains were reconstructed from a FIB-SEM acquisition of bloodstream-form trypanosomes. **(A)** A schematic illustration of the axonemal DMTs’ position relative to the PFR (magenta) and the cell body (CB). DMTs 7-8 face the CB, with DMT 7 marked by a longer connector to the PFR (asterisk). DMTs 3-4 and 7-8 are always positioned perpendicularly to the central pair (CP) of microtubules. **(B-D)** Representative FIB-SEM slices showing IFT trains (arrows) on DMTs 3-4 **(B)**, 7-8 **(C)**, or 1-2 **(D)**. The magenta asterisks indicate the longer PFR connector at the DMT7. Scale bar: 400 nm. **(E)** On the left, the 3D reconstruction of all flagella and IFT trains from a FIB-SEM acquisition. Each axoneme and its associated IFT trains are shown in different colours. The full volume is presented in **Video 2**. On the right, a representative flagellum is shown with IFT trains (red, arrowheads) localised on DMTs 3-4 and 7-8 of the axoneme (green). Scale bar: 900 nm. The 3D reconstruction of this flagellum is available in **Video 3**. **(F)** Distribution of IFT trains across axonemal DMTs (*n* = 224) in 29 flagella. **(G)** Length distribution (log scale) of IFT trains observed by FIB-SEM. **(H)** Length distribution of IFT trains across each set of DMTs.

Volumetric analysis of the IFT trains allowed for length measurement (*n* = 199) (**Fig. 2 G**). The number of analysed IFT trains was reduced due to the exclusion of incomplete ones from the FIB-SEM volume. In procyclic-form cells, IFT trains were categorised as short (∼ 200 nm) and long trains (∼ 900 nm) (Bertiaux et al., 2018). In bloodstream-form cells, we observed a similar distribution in length, where trains ranged from 100 nm to 6.2 µm, with no statistical distinction between short and long trains (p<0.0001; see Materials and Methods for information on the statistical tests used). On average, trains on DMTs 3-4 measured 1.03 ± 1.11 µm in length, while those on DMTs 7-8 measured 532 ± 515 nm. In procyclic-form trypanosomes, long trains averaged 822 nm on DMTs 3-4 and 968 nm on DMTs 7-8, with trains longer than 1.6 µm being rare (only one detected train with 2.5 µm of length). Conversely, bloodstream-form trypanosomes exhibited significantly longer IFT trains, reaching up to 6.2 µm, most of them found on DMTs 3-4, as indicated by the higher average length of the trains in these microtubules (**Fig. 2 H**). These data suggest that IFT functions similarly in both the bloodstream and procyclic life cycle stages, remaining restricted to DMTs 3-4 and 7-8.

### Multiple densities are detected in the proximal portion of the flagellum

In the reconstruction of the entire flagella length from the FIB-SEM series, some IFT trains (*n* = 12) appeared outside the DMTs 3-4 and 7-8 (**Fig. 2 F**). These trains were shorter than the others, typically ranging from 150 to 450 nm, with a maximum length of 1 µm (**Fig. 2 H**). Notably, eight of them were located at the proximal portion of the axoneme near the flagellar pocket. This prompted a detailed examination of the proximal portion of the flagellum in the FIB-SEM volume. We found eleven cells (10 monoflagellated and one biflagellated cells) where the proximal portion of the flagellum – between the basal body and the flagellar pocket exit – was visible (**Fig. 3, S2**). Remarkably, IFT trains outside the DMTs 3-4 and 7-8 were observed in four of these nine proximal segments.

**Figure 3:**
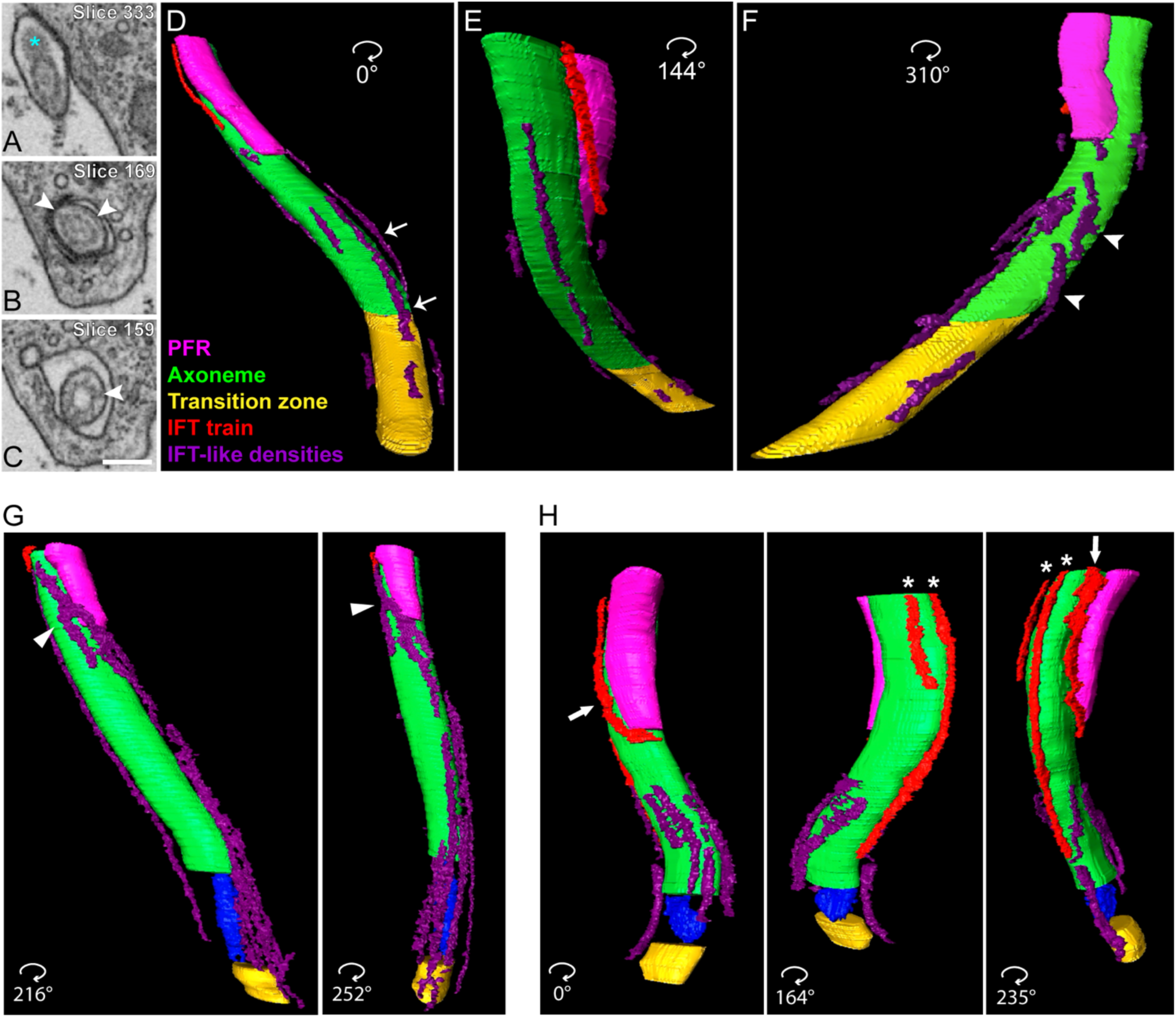
Densities surround all doublet microtubules of the flagellum portion inside the flagellar pocket. After reconstructing all proximal portions of flagella within the flagellar pocket in the same FIB-SEM series, as shown in Fig. 2, IFT-like densities were identified and annotated. **(A-C)** FIB-SEM slices of the same flagellum at different levels. **(A)** A slice of the axoneme outside the flagellar pocket, where the PFR (asterisk) is associated with DMTs 4- 7. **(B)** A slice of the axoneme inside the flagellar pocket, where the PFR is absent. Some IFT-like densities can be observed at this level (arrowheads). **(C)** A slice at the transition zone level, where another IFT-like density is visible (arrowhead). **(D-F)** The 3D reconstruction of the same flagellum shown in **(A-C)**. **(D)** A view of the reconstructed flagellum shows an IFT train (red) at DMTs 3-4, also revealing IFT-like densities (purple) at the level of the transition zone (yellow) and along the axoneme (green) portion without PFR (magenta). These densities are found on most DMTs, including those where the PFR will attach at a more distal level of the axoneme (arrows). **(E)** Another view of the same flagellum shows IFT-like densities on the DMTs opposite the side with the PFR. **(F)** A third view of the same flagellum confirms the presence of IFT-like densities on the DMTs that will be associated with the PFR. Some densities are found forming branches across multiple DMTs (arrowheads). The 3D reconstruction of this flagellum is shown in **Video 4**. **(G)** Two different views of the 3D reconstruction of another cell where a density seems to curve around the PFR (arrowheads) at the proximal portion of the flagellum. The full 3D reconstruction is shown in **Video 5**. **(H)** Three different views of the proximal portion of the flagellum from another cell reconstructed from the same FIB-SEM volume. This flagellum shows various IFT trains localised outside the DMTs 3-4 and 7-8 (asterisks), including one (arrows) that curves around the PFR at its initiation point. The full 3D reconstruction is displayed in **Video 6**. Green = axoneme, magenta = PFR, yellow = transition zone, blue = basal plate, red = IFT trains, purple = IFT-like densities.

Across all proximal segments, electron-dense material was consistently found between the axoneme and the flagellar membrane (**Fig. 3 A-C**, arrowheads). These materials followed the criteria for identifying IFT trains: located between the axonemal microtubules and the flagellar membrane, present in three or more consecutive slices. However, these densities exhibited thinner profiles and lower electron density compared to typical IFT trains. Upon reconstruction, these densities displayed profiles resembling IFT trains aligned along the microtubules (**Fig. 3 D-G, S2; Video 4**). However, they exhibited unique features, such as the presence of branches and regularly extending across multiple DMTs (**Fig. 3 F**, arrowheads). More specifically, when examining the distribution of these densities at the emergence of the PFR, they seem to be turning around the PFR (**Fig. 3 G**, arrowheads; **Video 5**) in four flagella, and at least one typical IFT train exhibited the same profile by extending across different DMTs (**Fig. 3 H**, arrow; **Video 6**).

After reconstructing all proximal flagellar portions, we assessed the DMTs for the presence of densities from the transition zone to the axoneme exit of the flagellar pocket.

Densities were detected across every DMT, including those to which the PFR would attach following the flagellum’s exit from the flagellar pocket (**Fig. 3 D**, arrows). Although FIB-SEM analysis does not permit the molecular identification of these densities, their profiles in the proximal portion of the axoneme resemble the IFT172 signal observed by 3D-STORM (**Fig. 1 B**, arrowhead). This resemblance suggests that these densities might correspond to anterograde IFT trains undergoing assembly and/or retrograde trains undergoing disassembly.

### Densities at the proximal portion of the flagellum contain IFT proteins

To characterise the IFT distribution at the proximal portion of the flagellum, where the densities were identified, we employed ultrastructure expansion microscopy (U-ExM). This technique offers improved resolution compared to conventional immunostaining fluorescence microscopy and allows molecular identification through antibody staining (Gambarotto et al., 2019). Bloodstream-form trypanosomes were expanded (with an expansion factor of approximately 6x based on the axoneme’s diameter) and stained with antibodies against alpha-and beta-tubulin and IFT172 (**Fig. 4**). Consistent with the STORM data, IFT172 concentrates at the proximal part of the transition zone (**Fig. 4 A**, green arrowheads), making a ring around the base of transition zone marker FTZC (Bringaud et al., 2000) (**Fig. 4 A**, white arrowheads; **Video 7**).

**Figure 4:**
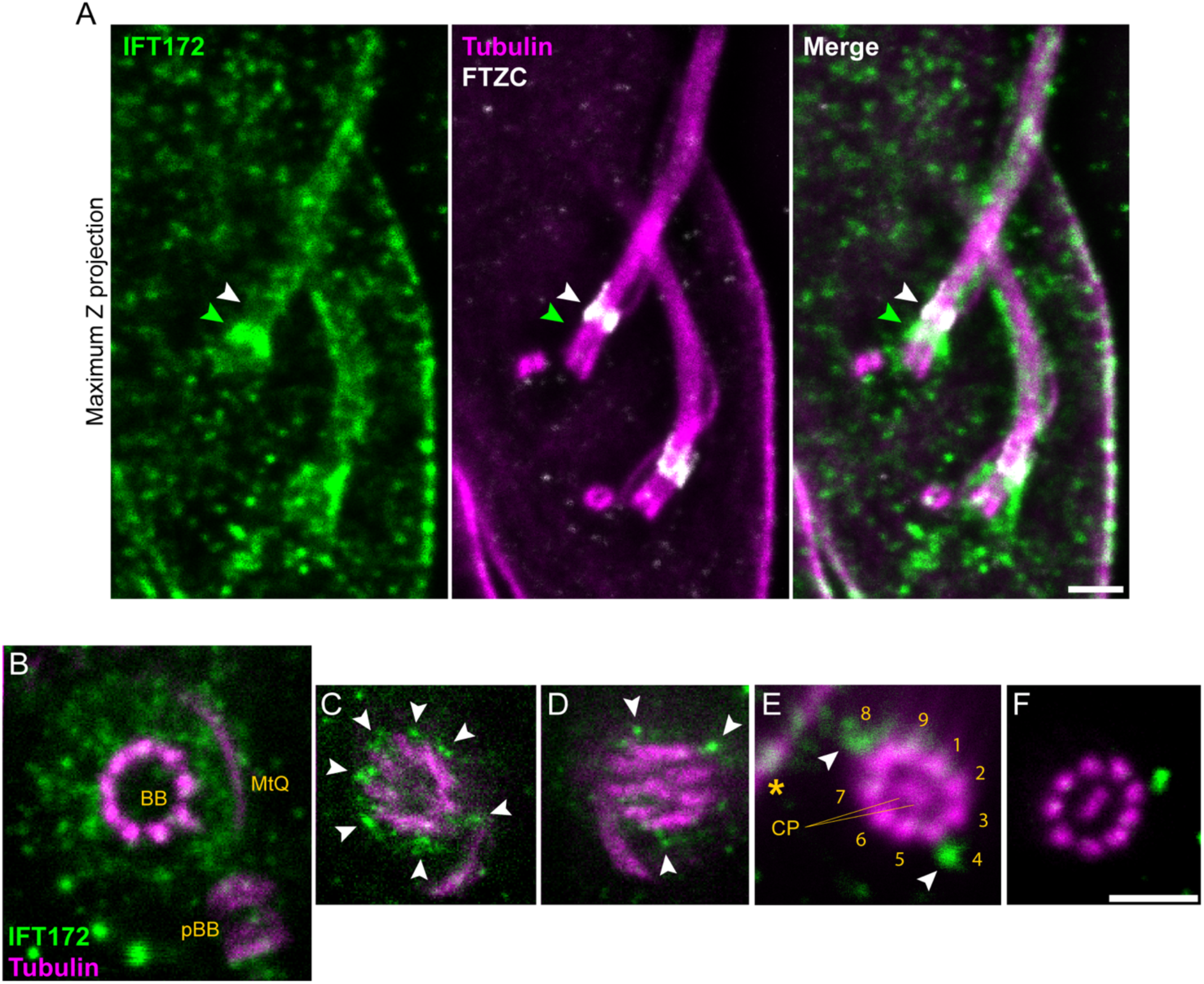
IFT translocates from nine to four doublet microtubules of the axoneme at the flagellar pocket level. Bloodstream-form trypanosome cells were expanded and stained with antibodies against alpha-and beta-tubulin (magenta) and IFT172 (green). **(A)** Maximum Z-projection confocal image of an expanded bloodstream-form trypanosome with two flagella, showing the IFT172 signal (green) at the proximal portion of the flagella (green arrowheads), marked by FTZC staining (white) of the transition zone (white arrowhead). The complete confocal volume is available in **Video 7**. Scale bar: 2 µm. **(B-F)** STED images of flagella from expanded bloodstream-form trypanosomes. **(B)** Optical slice showing pBB and BB with IFT172 signal extending from BB microtubules. The microtubule quartet (MtQ) is visible. **(C)** A more distal optical slice from the same cell as in **(B)** at the transition zone level, as indicated by the absence of the central pair, reveals the IFT172 signal in particles around most doublet microtubules (arrowheads). **(D)** A subsequent optical slice from the same cell in **(B-C)** reveals the central pair, with the IFT172 signal distributed in small particles along multiple doublet microtubules (arrowheads). **(E)** Image of another cell’s axoneme outside the flagellar pocket, showing subpellicular microtubules (yellow asterisk) of its cell body. The DMTs were numbered using the conventional references as described in Fig. 2 **B**, and the IFT172 signal was observed at DMTs 4 and 8 (arrowheads). **(F)** Optical slice of an axoneme at the level of the free tip showing IFT172 signal within one of the two tracks. As the cell body reference is missing, the DMTs could not be positioned. Scale bar: 1 µm.

To further enhance resolution, we combined U-ExM with a super-resolution technique called STimulated Emission Depletion (STED) microscopy (Hell and Wichmann, 1994) (**Fig. 4 B-F, S3**). The gels prepared for STED imaging were sliced and flipped so that the sectioned plane faced upward, allowing for better orientation during observation and improving our ability to resolve the stained elements. This approach enabled us to achieve sufficient resolution to individually observe the nine doublet microtubules and the central pair when present, facilitating precise localisation of the IFT172 signal along the flagellum.

At the basal body level, the IFT172 signal appeared weak and dispersed (**Fig. S3 A**). At the IFT basal pool level, IFT172 formed projections surrounding all nine DMTs, resembling the transition fibres (**Fig. 4 B, S3 B and C**). Moving distally of the same flagellum to the transition zone level (marked by the absence of the central pair of microtubules), these projections appear as more concentrated particles on the surface of most DMTs (**Fig. 4 C**, arrowheads). Further towards the tip, at the axoneme’s flagellar pocket level, the IFT172 signal manifested as small particles that remained present on many DMTs but were more restricted (**Fig. 4 D**, arrowheads).

The enhanced resolution allowed us to differentiate individual DMTs, which we numbered according to references used in the FIB-SEM analysis. In this numbering scheme, DMTs 7-8 always face the cell body, identifiable by the tubulin staining of the subpellicular microtubules (**Fig. 4 E**, asterisk). Also, the central pair is always parallel to the PFR and perpendicular to the DMTs 3-4 and 7-8. Last, the DMTs are numbered clockwise when the axoneme is viewed from the base of the flagellum (Lacomble et al., 2009). When the axoneme is outside the flagellar pocket (**Fig. 4 E and F, S3 D**), the IFT172 signal appears as stronger stained particles now sitting on opposite sides of the axoneme and positioned laterally relative to the central pair, corresponding to DMTs 3-4 and 7-8. In **Fig. 4 F**, which shows the axoneme at the free tip (marked by the absence of a cell body nearby), the DMTs can be easily distinguished, with an IFT172 particle positioned on one of the two IFT tracks. These observations support that the densities identified by FIB-SEM correspond to IFT material. They further indicate that at the flagellar pocket level, IFT trains are smaller and less organised, strengthening the hypothesis that this region is where IFT trains undergo assembly, disassembly, or both.

## Discussion

Among all species studied for intraflagellar transport (IFT), the positioning and distribution of IFT along the axoneme have been characterised in only two: *Chlamydomonas reinhardtii* and *Trypanosoma brucei*. In *Chlamydomonas*, IFT proteins are concentrated at the base of the flagellum, surrounding all nine doublet microtubules (DMTs), with mature IFT trains present on most of them (Jordan and Pigino, 2021). In contrast, *T. brucei* exhibits IFT trains on only four of the nine DMTs (Bertiaux et al., 2018). Little is known about the conservation of IFT positioning across ciliated organisms, and concluding based on two model organisms is premature. In this study, we clarified how IFT restriction occurs in trypanosomes.

The two hypotheses for explaining IFT restriction in trypanosomes were: 1) the restriction occurs at the basal pool, where IFT proteins would concentrate exclusively at the base of specific DMTs, or 2) the restriction would take place along the IFT tracks themselves, with IFT assembly limited to the four DMTs where the trains are present (Mallet and Bastin, 2019). Our data refute both hypotheses and instead support a third model. In this new model, IFT transitions progressively from a basal pool encircling all DMTs to IFT-like densities along the proximal portion of the flagellum and ultimately to IFT trains associated with only four DMTs. **Fig. 5** illustrates this proposed model.

**Figure 5:**
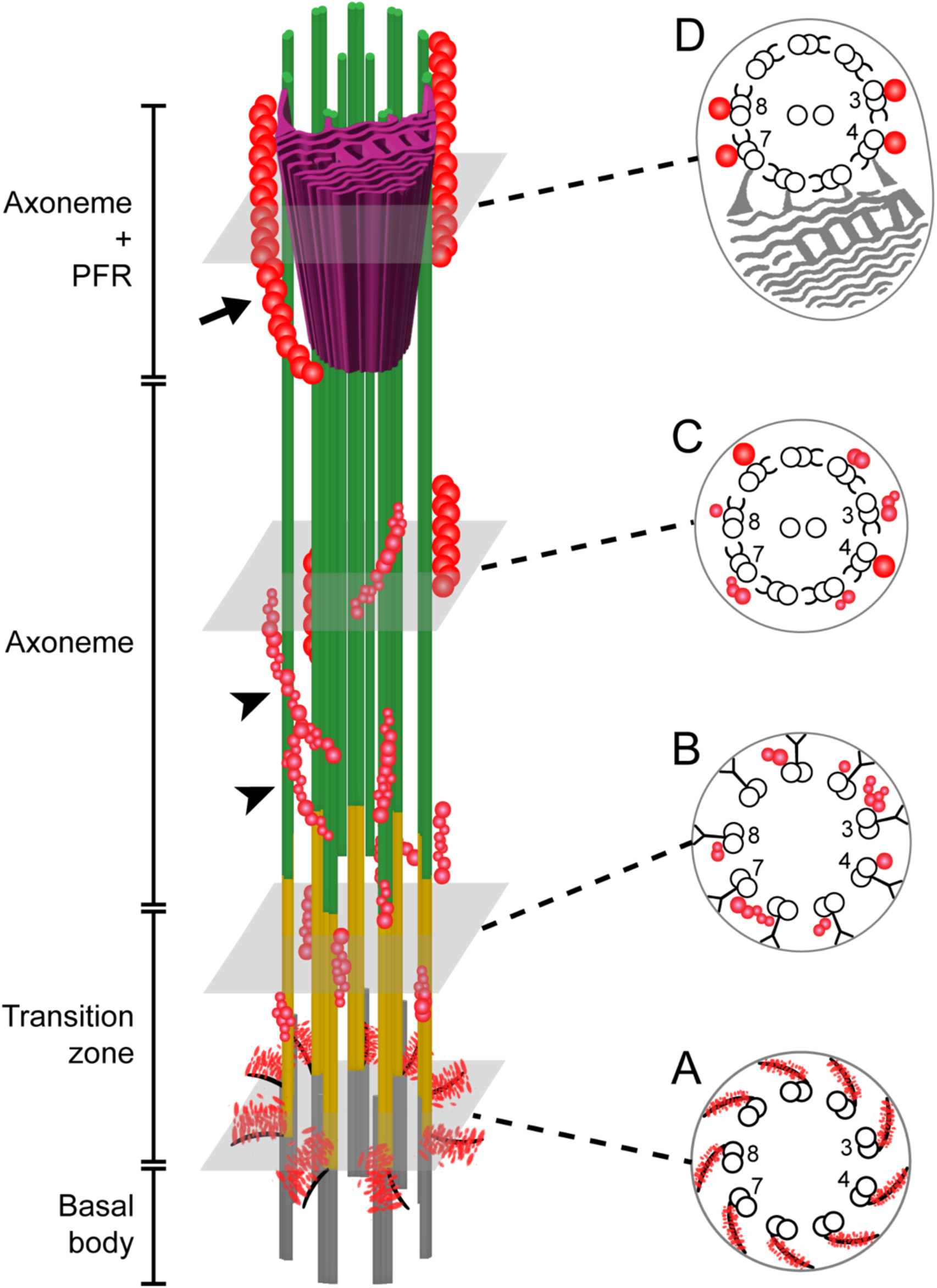
Proposed model for IFT restriction in *Trypanosoma brucei*. **(A)** IFT proteins (red) are concentrated above the transition fibres and surround all DMTs at the proximal portion of the transition zone (yellow). **(B)** In the transition zone, IFT-like densities are observed without restriction to specific DMTs. **(C)** As the axoneme (green) emerges from the transition zone, IFT-like densities and mature IFT trains appear associated with the DMTs. IFT-like densities often form branches atop multiple DMTs (arrowheads). At this level, IFT trains are also found outside the two usual sets of doublet microtubules. **(D)** When the axoneme exits the flagellar pocket, and the PFR (magenta) appears, IFT-like densities are no longer visible, and mature IFT trains are now only associated with DMTs 3-4 and 7-8. At the point where the PFR emerges, IFT trains seem to turn around it (arrow).

The first hypothesis was dismissed by the observation that IFT172 forms a nine-fold symmetrical pattern around all DMTs at the proximal portion of the transition zone, where the basal pool is located (**Fig. 5 A**). These projections likely represent IFT proteins associated with the transition fibres, as indicated by their partial colocalisation with the transition fibre marker RP2. This pattern is consistent with findings in *Chlamydomonas*, where immunocytochemistry demonstrated that IFT52 associates with the transition fibres (Deane et al., 2001). This result was further supported by immunofluorescence, which localised IFT-B core components, such as IFT81/74 and IFT172, as well as the IFT kinesin-II subunit FLA10 and the IFT-A component IFT139, to the distal end of the basal bodies, where transition fibres are situated (Richey and Qin, 2012). Additionally, a study using super-resolution structured-illumination microscopy (SIM) in *Chlamydomonas* showed that IFT-B proteins, including IFT57 and IFT172, along with IFT139, form a ring-like structure distal to the basal bodies (Picariello et al., 2019). Similarly, in RPE1 cells, super-resolution direct stochastic optical reconstruction microscopy (dSTORM) identified IFT88 with a nine-fold pattern at the transition fibres (Yang et al., 2018). In *C. elegans*, the IFT basal pool localises around the transition fibre protein FBF1, which is essential for the entry of IFT proteins into the cilium (Wei et al., 2013). Together, these observations support the idea that trypanosomes conserve the basal pool organisation of IFT proteins seen in other model organisms.

The second hypothesis, suggesting that IFT restriction occurs immediately after IFT proteins exit the basal pool, was refuted by our identification of IFT-like densities across all DMTs at the transition zone (**Fig. 5 B**). These densities were previously observed in conventional TEM as electron-dense material resembling IFT trains in the proximal portions of the flagellum (Absalon et al., 2008; Buisson, 2010; Subota et al., 2014). These IFT-like densities were examined in three dimensions, revealing their branching structures along multiple DMTs. Expansion microscopy confirmed the presence of IFT proteins with similar distribution. We propose that these IFT-like densities represent IFT trains undergoing assembly and/or disassembly. The observed branching (**Fig. 5**, arrowheads) could correspond to the fusion of IFT subcomplex polymers to form anterograde IFT trains or the fission of retrograde trains into IFT-A or IFT-B subcomplex polymers. This hypothesis aligns with the IFT assembly process described in *Chlamydomonas*, where the IFT-A and IFT-B proteins occupy distinct regions of the basal pool (Hou et al., 2007; Wingfield et al., 2017), reflecting the different contents of IFT-A and-B complexes along assembling IFT trains (van den Hoek et al., 2022). Given that these IFT-like densities are thinner and less electron-dense than typical IFT trains, we also propose that they may not be fully compacted due to the absence of associated motors. For example, IFT dynein, carried as cargo by anterograde trains, connects with both IFT-A and IFT-B complexes and may induce the final stages of anterograde train polymerisation. Silencing the expression of any component of the IFT dynein led to the formation of very short flagella, supporting the importance of IFT dynein in anterograde IFT trains (Blisnick et al., 2014). Similarly, the association of IFT kinesin could trigger the final steps of train assembly. In *C. elegans*, these final assembly steps occur at the proximal portion of the transition zone upon kinesin-II association with IFT trains (Mitra et al., 2024). In contrast, we did not observe typical IFT trains in the transition zone of *T. brucei*, suggesting that only the early stages of IFT assembly and/or the final stages of IFT disassembly take place in this region. Our approaches do not allow for distinct directionality of the IFT trains. In the literature, retrograde trains could not be detected at the ciliary base of *Chlamydomonas* (van den Hoek et al., 2022).

Strikingly, IFT-like densities were frequently observed in association with multiple DMTs, suggesting dynamic behaviour. Additionally, typical IFT trains were found on two DMTs near the exit of the flagellar pocket, appearing to turn around the PFR (**Fig. 5**, arrow), which suggests that IFT may switch between different microtubules in trypanosomes. Unfortunately, as the flagellar pocket is inside the cell body, its orientation makes it difficult to achieve enough resolution to resolve the IFT dynamics at the base by live imaging. As a result, it is currently not possible to evaluate the motility of these structures.

The final steps of IFT train assembly and/or the first steps of IFT train disassembly appear to occur at the flagellar pocket level, where both IFT-like densities and typical IFT trains are observed in association with the axoneme (**Fig. 5 C**). Typical IFT trains could be distinguished from IFT-like densities by their greater thickness, higher electron density, and lack of branching. At this stage, the PFR is not yet attached to the axoneme. However, once the PFR connects to DMTs 4-7, IFT trains are exclusively localised to DMTs 3-4 and 7-8 (**Fig. 5 D**), suggesting that the PFR may play a role in limiting IFT distribution. This hypothesis is further supported by the observation that both IFT-like densities and IFT trains frequently appear to turn around the PFR near the flagellar pocket exit. However, although the PFR might explain the absence of IFT on DMTs 5-6, due to the bridging connections between the PFR and the axoneme, IFT is also rarely observed on DMTs 1-2-9, which are not associated with the PFR. Thus, although these findings suggest a mechanism behind IFT selectivity, it is unlikely that the PFR is the sole factor responsible for IFT restriction.

Post-translational modifications of tubulin C-terminal tails may influence IFT train selectivity along specific DMTs (Mallet and Bastin, 2022). In *Spermatozopsis similis*, microtubules in the transition zone do not react with GT335, an antibody against the branching point of glutamate chains, indicating little or no polyglutamylation in this region compared to the axoneme (Lechtreck and Geimer, 2000). Variations in post-translational modifications along the flagellum could be detected by IFT molecular motors, guiding the engagement of the IFT trains with the microtubules (Janke and Magiera, 2020). This is supported by studies in mouse fibroblasts, where rapid axonemal deglutamylation led to slower anterograde IFT trains but not retrograde ones, resulting in the accumulation of IFT kinesin at the base of sensory cilia (Hong et al., 2018). The homodimeric kinesin-2 in humans (KIF17) and *C. elegans* (OSM-3) displayed increased speed and processivity on microtubules containing tyrosinated and polyglutamylated tubulin *in vitro* (Sirajuddin et al., 2014). Moreover, in the *C. elegans ccpp-1* mutant, which lacks an enzyme responsible for tubulin deglutamylation, the velocity of homodimeric IFT kinesin increased (O’Hagan et al., 2011). Collectively, these findings suggest that IFT kinesin processivity depends on DMT polyglutamylation at some level. In *T. brucei*, variations in axonemal polyglutamylation along the flagellum could affect kinesin engagement with DMTs at the flagellar pocket level. If DMTs 3-4 and 7-8 exhibit higher polyglutamylation levels, this could act synergistically with the PFR to modulate IFT train selectivity by influencing kinesin interaction with microtubules during train assembly.

Altogether, our findings suggest that IFT restriction in *T. brucei* arises from a combination of factors, which may include axonemal architecture, such as the presence of the PFR and motor protein interactions with the microtubules. These results provide new insights into the dynamics underlying selective IFT train localisation and reveal an unexpected ability of assembling (or disassembling) IFT trains to switch from one DMT to the other.

## Materials and methods

### Trypanosome culture and cell lines

The procyclic form of *T. brucei* strain Lister 427 was maintained in SDM79 medium supplemented with 10% FBS at 27°C. For N-terminal endogenous tagging of RP2 with mCherry, the first 497 nucleotides of RP2 (Tb927.10.14010) were chemically synthesised by GeneCust Europe for in-frame cloning using the HindIII and ApaI restriction sites of the p2845 vector (Kelly et al., 2007). The plasmid was linearised with Eco811 for integration in the *RP2* locus, ensuring *in situ* expression under the control of the 3’ untranslated region (UTR). Cells were nucleofected with the linearised plasmid (Burkard et al., 2007) and selected in a growth medium containing 1 μg/mL puromycin.

The bloodstream form of *T. brucei* strain Lister 427 was maintained in HMI-9 medium supplemented with 10% FBS at 37°C with 5% CO_2_. For N-terminal tagging of IFT81, 30 million cells were nucleofected with 10 µg of the p2675-mNG-IFT81 vector, which contains a puromycin resistance gene and was previously linearised with XcmI (Bertiaux et al., 2018). Nucleofection was performed using the Amaxa Nucleofector II system with program X-001, followed by selection in a growth medium containing 0.1 μg/mL puromycin.

### Super-resolution imaging by direct optical nanoscopy with axially localised detection (3D-STORM)

Procyclic-form trypanosome cells expressing mCh::RP2 were fixed and labelled for dSTORM imaging using antibodies against IFT172 (Absalon et al., 2008), revealed with an Alexa Fluor 647-coupled secondary antibody against mouse IgG1 diluted 1:500 (#A21237; Molecular Probes), and an antibody against dsRed that detects mCherry, revealed with an Alexa Fluor 555-coupled secondary antibody against rabbit antibodies diluted 1:500 (#A27039; Molecular Probes). 2×10^7^ cells per round 25 mm coverslip were harvested and fixed for 5 minutes in 4% formaldehyde in PBS, followed by 5 minutes in-20°C methanol and 15 minutes rehydration in PBS. Primary and secondary antibodies were incubated for 45 minutes at 37°C in a humid environment. After the final wash, cells were post-fixed with 4% PFA for 5 minutes, followed by two additional washes in PBS. Samples were stored in PBS until acquisition. For dual colour imaging, a sequential strategy was adopted, imaging first the AF647-labelled proteins and afterwards the AF555-labelled proteins. Labelled cells were incubated in dSTORM buffer (SMART kit, Abbelight) and immediately imaged on an Abbelight SAFe360 module in direct optical nanoscopy with axial localised detection (DONALD) configuration (Bourg et al., 2015; Deschamps et al., 2014). This enables the detection of single fluorophores with a lateral localisation precision of typically 5 to 10 nm and an absolute vertical position with a precision of 15 nm, and it is optimally adapted for 3D super-resolution imaging of biological structures within 500nm of the coverslip. The dual view super-resolution module SAFe360 (Abbelight, Cachan) was equipped with two Hamamatsu Orca FLASH4 sCMOS cameras and mounted on an Olympus IX83 inverted microscope with a 1.49NA 100X oil objective and ZDC focus control; the sample was excited in HiLO using a 640 nm laser (for AF647) and a 532 nm laser (for AF555), with a Starscan azimuthal TIRF (Errol). Once the optimal photo-switching was established, in DONALD configuration, two simultaneous acquisition films were collected (50 ms per frame, 50000 frames per acquisition) and used to perform lateral and axial localisation of individual fluorophores with Abbelight NEO software. Drift correction, data post-processing and visualisation were also performed using NEO software.

### BSF live cell imaging

*T. brucei* AnTat1.1 bloodstream-form cells expressing mNG::IFT81 were imaged using a DMI 4000 B microscope (Leica) and an oil-immersion objective Plan Apochromat HCX 100x/1.4 NA (Leica), which was equipped with an Evolve 512 EMCCD camera (Photometrics). Cells were incubated at 20°C for 10 minutes in a CherryTemp temperature-controlling chamber (Cherry Biotech), which caused them to stop swimming, enabling the imaging of mNG::IFT81 particles. Following 30-second acquisitions with 100 ms exposure time, kymographs were generated using the Icy software (de Chaumont et al., 2012).

### TEM

*T. brucei* AnTat1.1 bloodstream-form cells were fixed overnight at 4°C in a solution containing 2.5% glutaraldehyde (#16210; Electron Microscopy Sciences) and 2% formaldehyde (#15714; Electron Microscopy Sciences) in 0.1 M sodium cacodylate buffer (pH 7.2). After fixation, the samples were washed three times with 0.1 M sodium cacodylate buffer and three times with milli-Q^®^ (Merck) water. The samples were then incubated with 2% uranyl acetate (#22400; Electron Microscopy Sciences) in milli-Q water for 1 hour at room temperature, followed by three washes with milli-Q water. Dehydration was performed in a graded ethanol series of 25%, 50%, 75%, and 95% ethanol in milli-Q water for 10 minutes each, followed by three washes in 100% ethanol. The samples were then incubated for 15 minutes in propylene oxide and embedded overnight in a 1:1 solution of propylene oxide and the epoxy resin Agar 100 (#AGR1031; Agar Scientific). This was followed by three 1-hour incubations in pure epoxy resin. The samples were placed in a silicon mould with epoxy resin and polymerised at 60°C for 48 hours. 50-70 nm sections were obtained using an EM UC6 ultra-microtome (Leica) and stained with 2% uranyl acetate in milli-Q water and 80 mM lead citrate in milli-Q water before imaging. The sample was imaged using a Tecnai BioTWIN 120 microscope (FEI) equipped with a MegaView II camera (Arecont Vision).

### FIB-SEM, data segmentation and length measurements

The FIB-SEM series of *T. brucei* AnTat1.1 bloodstream-form trypanosomes was acquired previously (Vanwalleghem et al., 2015). Following image acquisition, the image stack was aligned using ImageJ software, and subsequent manual segmentation and 3D reconstructions were performed with Amira software (ThermoFisher). Each electron-dense particle located between the axoneme and the flagellar membrane that appeared in three or more consecutive slices was annotated as an IFT train (Bertiaux et al., 2018). To identify IFT-like densities, we applied the same criteria as for IFT trains.

Amira 2021.2 software was utilised for length measurements. Initially, the objects were skeletonised using the Centerline Tree module, followed by the Spatial Graph View module to generate filaments corresponding to the segmented objects. Finally, the Spatial Graph Statistics module was used to obtain the length of the filaments. The filaments were manually identified to correspond with the segmented objects.

The GraphPad Prism 10 software was used to test for the normality (Gaussian) distribution of the IFT trains’ length into two populations. The tests used were D’Agostino & Pearson, Anderson-Darling and Shapiro-Wilk.

### Expansion Microscopy

10^6^ *T. brucei* bloodstream-form cells were washed once in trypanosome dilution buffer (TDB; 20 mM Na_2_HPO_4_, 2 mM NaH_2_PO_4_, 80 mM NaCl, 5 mM KCl, 1 mM MgSO_4_, 20 mM glucose, pH 7.4) and fixed with 4% formaldehyde in TDB at 37°C for 15 minutes. After two washes in PBS, cells were adhered to poly-L-lysine-covered 12 mm coverslips for 1 hour. The following steps were done as described by Louvel et al. (2023). The samples were post-fixed overnight in 0.1% (w/v) N, N′-(1,2-Dihydroxy-1,2-ethanediyl) bisacrylamide (DHEBA), 10% (w/v) acrylamide and 19% (w/v) sodium acrylate in PBS at room temperature. For the gelation, gelation chambers were placed on ice, the coverslips were transferred to the chambers and the sample coverslip was covered with monomer solution (10% acrylamide, 19% sodium acrylate, 0.1% DHEBA, 0.25% tetramethyl ethylenediamine (TEMED) and 0.25% ammonium persulfate). Samples were incubated for 15 minutes on ice, then at 37 °C for 45 minutes to finish the gelation. The coverslips were then removed from the gelation chamber and placed in the denaturation buffer (200 mM sodium dodecyl sulphate (SDS), 200 mM NaCl and 50 mM Tris-BASE, pH 6.8) until the gel detached from the coverslip. The gel was placed in 1.5 mL Eppendorf tubes containing 1 mL of the denaturation buffer and incubated at 85°C for 90 minutes. The gels were transferred to Petri dishes and expanded by three consecutive 30- minute incubations in milli-Q water (Louvel et al., 2023).

Samples were stained using antibodies against alpha-tubulin (#ABCD_AA345; ABCD Antibodies) and beta-tubulin (#ABCD_AA344; ABCD Antibodies) diluted 1:200, revealed with an Alexa Fluor 594-coupled secondary antibody against mouse IgG2a (#115-585-206; Jackson ImmunoResearch) diluted 1:400. Samples were co-stained with anti-IFT172 antibody (Absalon et al., 2008) diluted 1:50, revealed with an Alexa Fluor 488 (#115-545-205; Jackson ImmunoResearch) or ATTO 647N (#610-156-040; Rockland Immunochemicals) coupled secondary antibody against mouse IgG1 diluted 1:400. Anti-FTZC (Bringaud et al., 2000) was diluted 1:500, revealed with Alexa Fluor 488-coupled secondary antibody against rabbit (#211-542-171; Jackson ImmunoResearch) diluted 1:400. For staining, a gel piece of 2×2 cm was shrunk in PBS for 10 minutes, then incubated with the primary antibodies diluted in 2% BSA in PBS overnight at 37°C under agitation. After three washes of 10 minutes each in PBS-Tween (0.1% Tween 20 in PBS), the gel was incubated with the secondary antibodies diluted in 2% BSA in PBS for 4 hours at 37°C under agitation. The gel was then washed three times for 10 minutes each in PBS-Tween, followed by expansion in milli-Q water by three consecutive incubations of 30 minutes each. Before imaging, the gel was cut into cross-sectional slices approximately 2 mm thick using a blade. The slices were then flipped so that the sectioned plane faced upward, and they were placed in a µDish coated with poly-L-lysine and sealed with eco-sil two-component silicon (#1300 6100; Picodent). Images were acquired on a DMi SP8 confocal microscope (Leica) equipped with an oil-immersion objective Plan Apochromat CS2 63x/1.4 NA (ZEISS) or on a STimulated Emission Depletion microscope (STED) expert line microscope (Abberior Instruments) using a 100X/1.4 NA oil objective (Olympus) and the Avalanche Photo-Diodes (APDs, single point photon counting modules) detector (Excelitas). The two-colour STED images were obtained in line-interleaved mode, switching on one excitation laser line at a time, 561nm for Alexa 594 and 640 nm for ATTO 647N. The laser line for STED was 775 nm, pulsed. Pixel sizes were between 10 and 25 nm, and elementary pixel dwell times of 1 μs with up to a few tens of line averages, depending on the acquisitions (Abberior).

## Supplemental material

The supplemental material includes two figures and six videos. **Fig. S1** shows IFT in live bloodstream-form cells and IFT positioning using TEM. **Fig. S2** presents all the reconstructed flagella observed by FIB-SEM and the distribution of IFT-like densities. **Fig. S3** exhibits other STED images of IFT172 and tubulin staining in expanded trypanosomes. **Video 1** displays mNG::IFT81 particles moving along the flagella of the two cells shown in Fig. S1 A. **Video 2** features the complete FIB-SEM series of bloodstream-form trypanosomes and the 3D view of all reconstructed axonemes and IFT trains shown in Fig. 2 A. **Video 3** provides a 3D view of one reconstructed axoneme exhibiting IFT trains on DMTs 3-4 and 7-8, as displayed in Fig. 2 A. **Video 4** shows the 3D view of the reconstructed flagellum presented in Fig. 3 D-F. **Video 5** displays the 3D view of the reconstructed flagellum in Fig. 3 G. **Video 6** provides the 3D view of the reconstructed flagellum shown in Fig. 3 H. **Video 7** provides the confocal volume of the trypanosome shown in Fig. 4 A.

## Data Availability

## Acknowledgements

We thank D. Pérez-Morga (ULB, Belgium) for providing the FIB-SEM data set. We thank Vincent Louvel, Virginie Hamel and Paul Guichard (University of Geneva, Switzerland) for their contribution to the U-ExM experiments. We are grateful to the Photonic Bioimaging facilities for access to their equipment. We are thankful for the support of the UBI equipment from the French Government Programme Investissements d’Avenir France BioImaging (FBI, N° ANR-10-INSB-04-01).

This work was supported by La Fondation pour la Recherche Médicale (FDT20170436836), the ANR (ANR-18-CE13-0014-01) and a French Government Investissement d’Avenir programme, Laboratoire d’Excellence “Integrative Biology of Emerging Infectious Diseases” (ANR-10-LABX-62-IBEID). This work was also supported by *Région Ile-de-France* (DIM ELICIT) and *Institut Pasteur* (STED microscope).

Author contributions: A.A. Alves: Project administration, Data curation, Formal analysis, Investigation, Methodology, Resources, Validation, Visualisation, Writing - original draft, Writing - review & editing, J. Jung: Data curation, Formal analysis, Methodology, G. Moneron: Data curation, Formal analysis, Methodology, H. Vaucelle: Data curation, C. Fort: Data curation, J. Buisson: Data curation, C. Schietroma: Data curation, Formal analysis, P. Bastin: Conceptualization, Funding acquisition, Supervision, Writing - review & editing.

## Video legends

**Video 1: IFT in live bloodstream-form trypanosomes.** Video showing the displacement of mNG::IFT81 particles along the flagellum in the two cells presented in Fig. S1 A. Scale bar: 5 µm.

**Video 2: 3D view of the FIB-SEM volume with segmented axoneme and IFT trains (Fig. 2 A).** A 3D visualisation of the FIB-SEM volume where each axoneme and IFT train identified in the series has been segmented and displayed in distinct colours.

**Video 3: 3D view of the segmented axoneme and associated IFT trains (Fig. 2 A).** A 3D view of the segmented axoneme (green) with associated IFT trains (red) on DMTs 3-4 and 7-8, as shown in Fig. 2 A, right panel.

**Video 4: 3D view of the reconstructed flagellar pocket (Fig. 3 A).** A 3D visualisation of the reconstructed flagellar pocket showing the proximal portion of the axoneme, including the transition zone (yellow), the axoneme (green) within the flagellar pocket, and a segment of the axoneme exiting the flagellar pocket with the PFR (magenta) visible. A mature IFT train (red) and multiple IFT-like densities (purple) are also visible.

**Video 5: 3D view of the reconstructed flagellar pocket (Fig. 3 G).** A 3D view of another reconstructed flagellar proximal portion, with IFT-like densities (purple) detected along the transition zone (yellow) and the axoneme (green) within the flagellar pocket. One density looks like it is turning around the PFR (magenta) at its emergence point. The basal plate is shown in blue, and a small part of an IFT train is in red.

**Video 6: 3D view of the reconstructed flagellar pocket (Fig. 3 H).** A 3D view of another proximal portion of the flagellum, showing IFT trains (red) outside DMTs 3-4 and 7-8 and one that seems to turn around the PFR (magenta). IFT-like densities (purple) are also observed along the proximal portion of the axoneme (green) within the flagellar pocket. The basal plate is shown in blue.

**Video 7: Full confocal volume of a bloodstream-form cell (Fig. 4 A).** Z-stack of a bloodstream-form trypanosome containing two flagella stained with antibodies against tubulin (magenta), FTZC (white), and IFT172 (green).

## Notes

### Competing Interest Statement

The authors have declared no competing interest.

