## Supplementary figures 1-3 for "Intraflagellar Transport Selectivity Occurs with the Proximal Portion of the Trypanosome Flagellum"

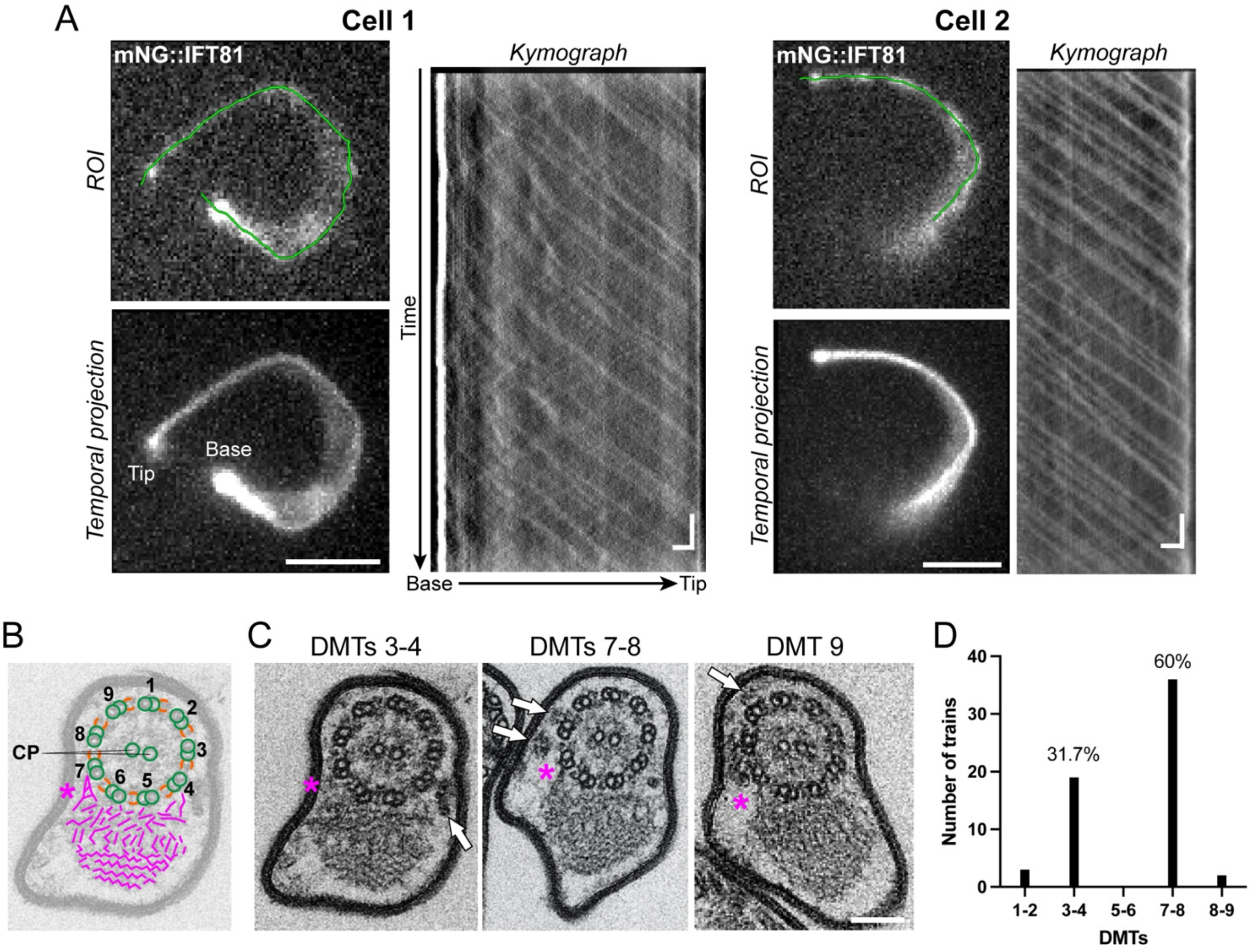


**Supplemental Figure 1: IFT in the bloodstream form of *Trypanosoma brucei*. (A)** Two representative cells expressing mNG::IFT81 were imaged, and particles moving along the flagellum were detected. A region of interest (ROI, green line) was drawn along the flagellum using the temporal projection to extract the kymographs from the acquisitions. The kymograph exhibits the particles' displacement from base to tip (anterograde transport) and from tip to the base (retrograde transport) of the flagellum. Scale bar: 5 µm. Kymograph horizontal scale bar: 2 µm; vertical scale bar: 2 s. The full acquisitions can be visualised in **Video 1**. **(B)** A schematic diagram illustrates that the DMTs (green) are numbered in a clockwise direction relative to the orientation of the dynein arms (orange). Notably, the DMT7 features a long and thick connection with the PFR (asterisk). **(C)** Representative TEM images show flagella with IFT trains localised on DMTs 3-4, 7-8 or 9 (arrows). The magenta asterisks highlight the long PFR connector associated with DMT7. Scale bar: 100 nm. **(D)** The distribution of IFT trains on axoneme DMTs (*n* = 60) was quantified across 41 transversal flagella sections obtained by TEM.

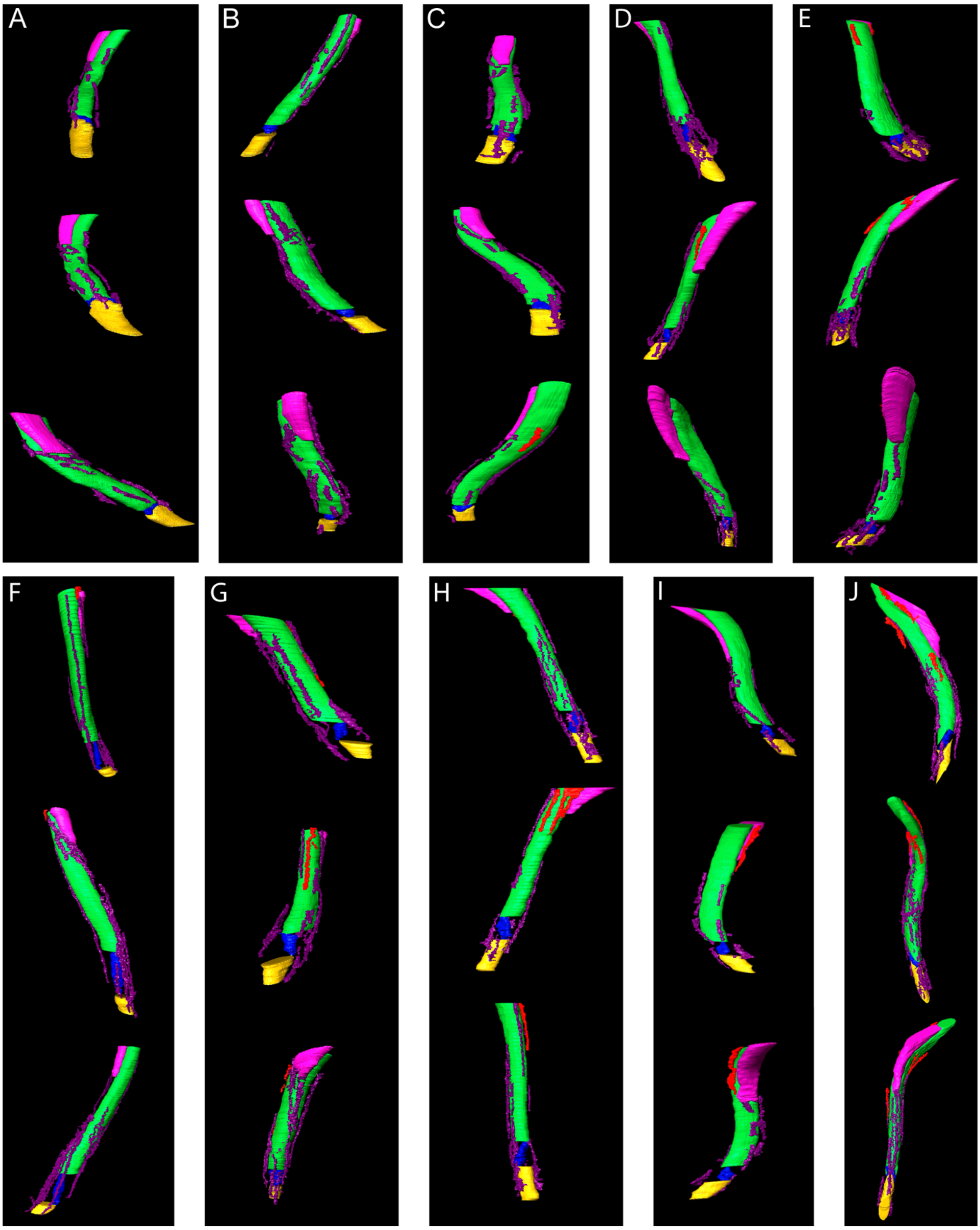


**Supplemental Figure 2: IFT-like densities along axoneme portions within the flagellar pocket (A-J).** All proximal portions of flagella within the flagellar pocket found in the bloodstream-form trypanosomes' FIB-SEM series were reconstructed, and IFT-like densities were observed in each of them. In the panel, the reconstructed flagella are shown with X-axis rotations to illustrate the presence of IFT-like densities surrounding the axoneme. Green = axoneme, magenta = PFR, yellow = transition zone, blue = basal plate, red = IFT trains, purple = IFT-like densities.

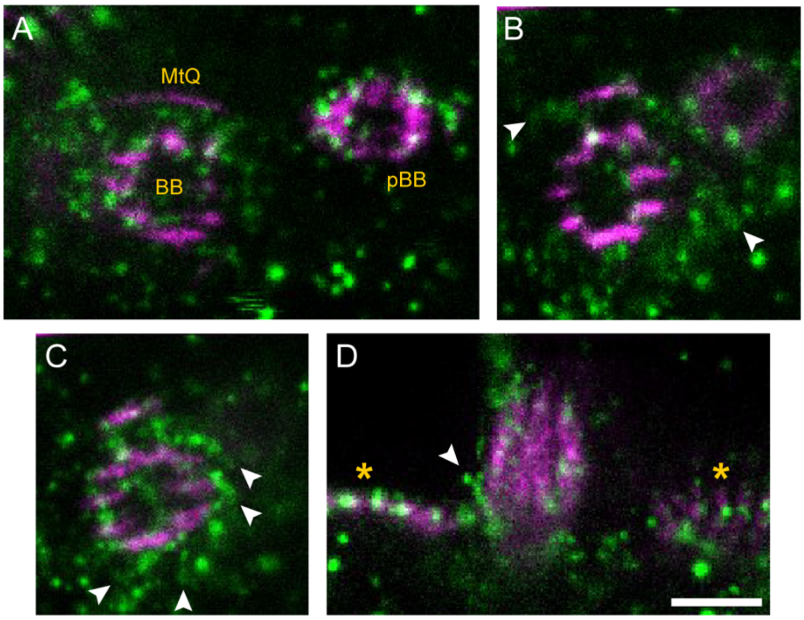


**Supplemental Figure 3: STED images of IFT172 (green) and tubulin (magenta) of expanded bloodstream-form trypanosomes. (A)** Axoneme at the basal body (BB) level showing a diffuse and weak IFT172 signal. pBB: pro-basal body, MtQ: microtubule quartet. **(B)** Axoneme at the very beginning of the transition zone starts exhibiting the IFT172 signal extending from some DMTs (arrowheads). **(C)** A subsequent optical slice from the same cell in **(B)** shows the flagellum at the proximal portion of the transition zone, showing IFT172 projecting from the DMTs (arrowheads). **(D)** An optical slice of an axoneme at the level of the flagellar pocket exit, marked by the subpellicular microtubules (yellow asterisks), shows an IFT train curving around the DMTs (arrowhead). Scale bar: 1 µm.
